# Whole-body Loss of FSP27 Impairs Cognitive Function via Disruption of Neuro-Metabolic Pathways

**DOI:** 10.1101/2025.07.28.667235

**Authors:** Andrew Pugh, Aarav Bhasin, Connor Aleshire, Rabia Basri, Chloe Becker, Ishika Puri, Hebaallaha Hussein, Kaia Mckinney, Kate Tenerowicz, Murali Vijayan, Bijinu Balakrishnan, Vishwajeet Puri

**Author notes:** Corresponding Author Vishwajeet Puri, M.S., Ph.D. **Author Contributions:** Equivalent first authors (Andrew Pugh, Aarav Bhasin).

## Abstract

Fat-Specific Protein 27 (FSP27), originally identified for its role in adipocyte lipid metabolism and energy homeostasis, plays a key role in regulating lipolysis and maintaining insulin sensitivity. Beyond adipose tissue, emerging studies have uncovered its involvement in hepatic and skeletal muscle function. More recently, FSP27 has also been implicated in maintaining vascular health through its influence on endothelial signaling. Despite growing insights into FSP27’s systemic functions, its involvement in the central nervous system and cognitive regulation have remained unexplored. In this study, we present the first evidence that FSP27 is a critical regulator of cognitive function. Utilizing a global FSP27 knockout (Fsp27^-/-^) mouse model, we demonstrate that FSP27 deficiency results in significant impairments in learning, memory retention, and spatial awareness. To elucidate the underlying molecular mechanisms, genome-wide transcriptome profiling of brain tissue from Fsp27^-/-^ mice was performed, revealing significant (P<0.05) alterations in gene expression related to neurocognition and metabolic pathways. Notably, FSP27 deletion was associated with genomic instability (P=0.05), downregulation of genes essential for axonal transport (NES=-1.91, FDR q-value=0.21), neuronal plasticity, and brain development (NES=-1.91, FDR q-value=0.28), along with signatures of disrupted systemic metabolism and elevated stress responses (NES=1.81, FDR q-value=0.20) in the brain, the processes tightly linked to cognitive dysfunction. Collectively, these findings establish FSP27 as a molecular node connecting metabolic regulation with cognitive health and identify it as a promising target for therapeutic intervention in neurodegenerative and cognitive disorders.

**Significance Statement:** FSP27, previously known for its role in lipid metabolism, is now revealed to be a critical regulator of brain function. Our study provides the first direct evidence that FSP27 supports learning and memory by maintaining neuronal integrity and regulating neurocognitive gene networks. These findings uncover an unexpected metabolic– cognitive link and position FSP27 as a promising molecular target for therapeutic strategies aimed at preventing or treating neurodegenerative and cognitive disorders.

## Introduction

Cognition encompasses a broad range of mental processes involved in acquiring, processing, storing, and retrieving information(1). These processes include perception, attention, memory, learning, and reasoning. Neurocognition, more specifically, focuses on how these cognitive functions are supported by underlying brain structures and neural mechanisms(2). As individuals age, cognitive functions naturally decline, often leading to impairments in memory, reasoning, and processing speed (3). In some cases, this decline can progress into neurodegenerative disorders such as Alzheimer’s disease, highlighting the importance of understanding molecular and physiological contributors to brain health.

The human brain is composed of approximately 60% fat, making it the most lipid-rich organ in the body (4). These fats are essential for constructing neuronal membranes and myelin sheaths, as well as maintaining synaptic integrity (4, 5). Disturbances in lipid metabolism can disrupt these functions, potentially impairing learning, memory, and other aspects of cognition. Since lipid homeostasis plays a central role in maintaining brain health, proteins involved in fat storage and utilization may have underrecognized impacts on cognitive function.

Studies have shown that the brain and skeletal muscle are interconnected, sharing molecular signaling pathways that impact cognition, mood, and neuroplasticity (6, 7). Physical activity, particularly muscle contraction, stimulates the release of myokines which travel through the bloodstream to the brain and exhibit neuroprotective effects. Among these, brain-derived neurotrophic factor (BDNF) is one of the most well-studied and is known to support synaptic plasticity, learning, and memory by promoting neuronal survival and growth(8, 9). Similarly, irisin, a cleaved product of the muscle-expressed protein FNDC5, has been shown to upregulate BDNF in the hippocampus and enhance neural differentiation and memory function (10, 11). These findings emphasize that metabolic health and muscle function can strongly influence brain performance, raising the possibility that disturbances in muscle-derived signaling may contribute to cognitive dysfunction.

Fat-Specific Protein 27 (FSP27), also known as cell death-inducing DFFA-like effector C (CIDEC), was originally identified as a key regulatory protein in fat metabolism in adipose tissue(12–18), where its expression is positively associated with glucose homeostasis in humans(14, 17) (19–21). Although the whole-body Fsp27^-/-^ mouse model showed higher oxidative metabolism with improved insulin sensitivity(15, 22), various tissue-specific studies including liver, adipose and endothelial cells, have unequivocally highlighted the importance of FSP27 protein in supporting metabolic health and promoting improved whole-body metabolism in mice(23–25). Our recent study in whole-body Fsp27^-^/-mice showed a drastic decline in muscular fat storage, muscle endurance, and muscle strength(26).

Given FSP27’s key roles in lipid regulation and energy balance in adipose tissue and muscle, both of which influence cognitive health(27–31), we hypothesized that FSP27 may also impact brain physiology and cognitive function. This study aimed to test this hypothesis by examining cognitive performance in whole-body Fsp27^-/-^ mice. To assess behavioral outcomes, we conducted a series of neurobehavioral tests, including the Morris Water Maze(32), Labyrinth Maze(33), and Y-Maze(34, 35) with wild-type (WT) and Fsp27^-/-^ mice. These tests were designed to evaluate learning capacity, memory, and spatial awareness. In parallel, we performed transcriptomic profiling via RNA sequencing on brain tissue from WT and Fsp27^-/-^ mice to identify gene expression and pathway alterations associated with FSP27 deletion.

## Results

### Fsp27^-/-^ Mice Demonstrated Diminished Learning Capabilities in the Morris Water Maze

To assess cognitive performance, the Morris Water Maze test was performed on wild-type (WT) and Fsp27^-/-^ male mice. As described in the methods, the mice were trained from day 1 to day 4 with the platform visible on day 1, but submerged on days 2-1 The final cognitive test in the Morris Water Maze was performed on day 5 with the platform removed. On day 1 of the training period, with a visible platform, no significant difference was observed in the latency to reach the platform between WT and Fsp27^-/-^ mice, indicating comparable visual and motor abilities at baseline (Fig. 1A). Furthermore, the average swimming speed was 0.25 m/s, indicating that there was no difference in the swimming capability of WT or Fsp27^-/-^ mice. On days 2-4, the platform was submerged to assess spatial learning. WT mice exhibited a rapid and progressive reduction in escape latency across training days, reflecting intact learning capabilities (Fig. 1B-D). In contrast, Fsp27^-/-^ mice showed minimal improvement over time, suggesting impaired spatial learning.

**Figure 1.**
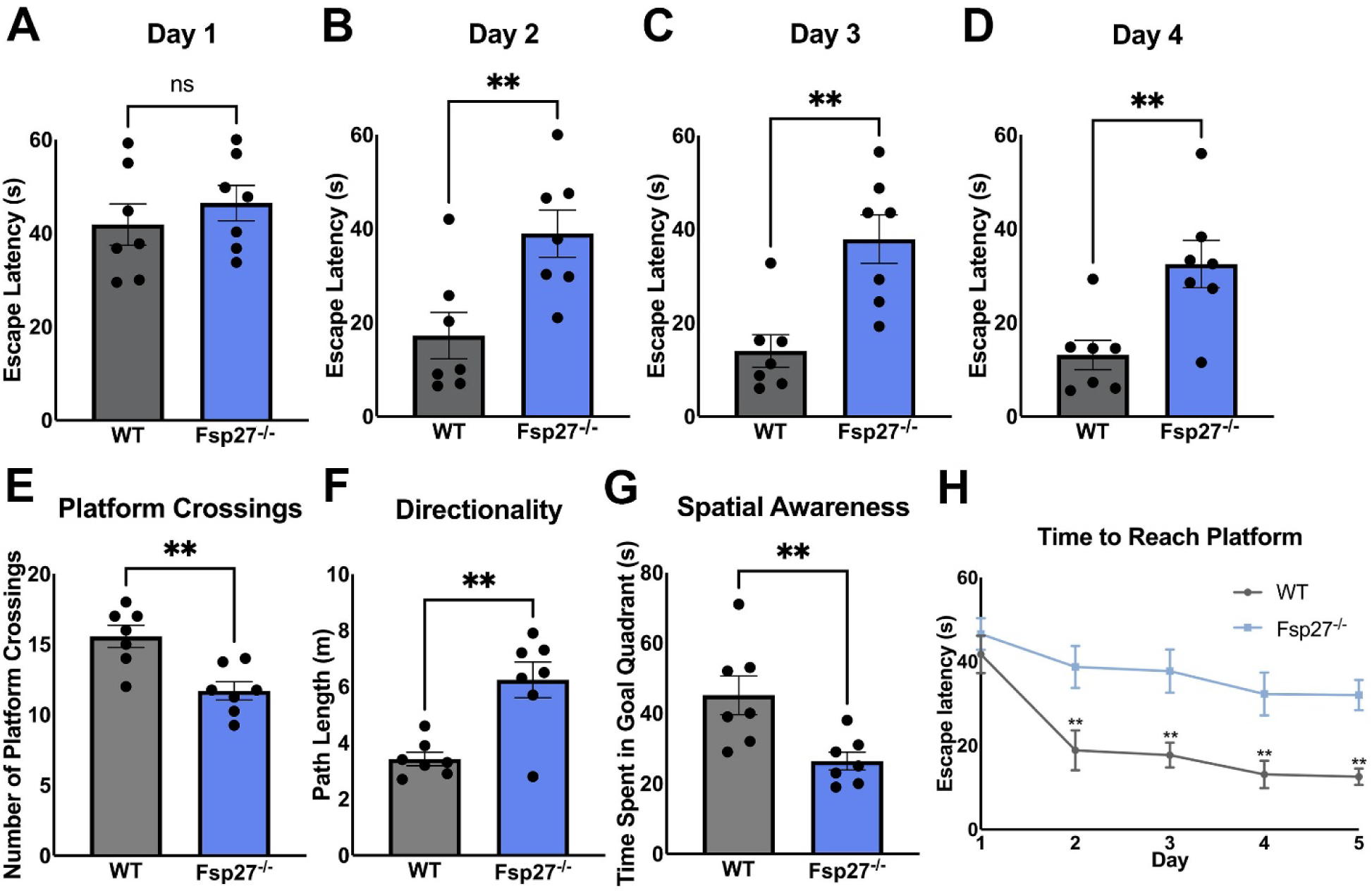
Fsp27^-/-^ mice demonstrated reduced learning capacity and spatial awareness. Time to reach out to the platform in the water maze on (A) day 1 (B) day 2 (C) day 3 and (D) day 4. *\*\*P* ≦ 0.01 ;Unpaired Student t-test. (E) Number of times the mouse approached within 1 cm of the 8 cm former platform location on day 5. (F) Path length to the previous platform location on day 5. (G) Time spent in the goal quadrant on day 5 (quadrant 1). (H) Time taken to reach the escape platform for WT (grey) and Fsp27KO (blue) mice across all five days. Data presented as mean ± SEM. *\*P* ≦ .05, *\*\*P* ≦ 0.01, *\*\*\*P* ≦ 0.001, *\*\*\*\*P* ≦ .0001; Unpaired Student t-test.

On day 5, the trial was conducted with the platform removed to assess spatial memory by measuring a mouse’s ability to recall the location of the previously learned platform. WT mice crossed over the former platform location more frequently (Fig. 1E), took a more direct path to the target (Fig. 1F), and spent significantly more time in the target quadrant (Fig. 1G). Conversely, Fsp27^-/-^ mice displayed disorganized swimming patterns, often circling the periphery of the pool, and taking longer path lengths to the target. These mice also made fewer crossings over the previous platform location and spent less time in the target quadrant, indicating a deficit in spatial memory. Overall, our results suggested a significantly slower learning process and impaired spatial memory in Fsp27^-/-^ mice compared to WT mice.

### Labyrinth Maze Confirms Deficit in Learning Capabilities in Fsp27^-/-^ Mice

To further confirm impaired learning capabilities in the Fsp27^-/-^ mice, the labyrinth maze test was performed. The same cohort of mice used in Fig. 1 were trained on the labyrinth maze for 2 days, followed by the final test day. As shown in Fig. 2, on day 1, there was no significant difference in latency to escape the maze between WT and Fsp27^-/-^ mice. However, on day 3 the Fsp27^-/-^ mice took significantly longer to escape the maze (Fig. 2). While the WT mice showed improvement in latency to escape the maze, Fsp27^-/-^ mice showed minimal to no progress indicating challenges with learning. These results confirmed impaired memory, learning, attention, and decision-making in Fsp27^-/-^ mice. These behaviors reflect poor spatial learning, working and reference memory, cognitive flexibility and problem-solving abilities in Fsp27^-/-^ mice.

**Figure 2.**
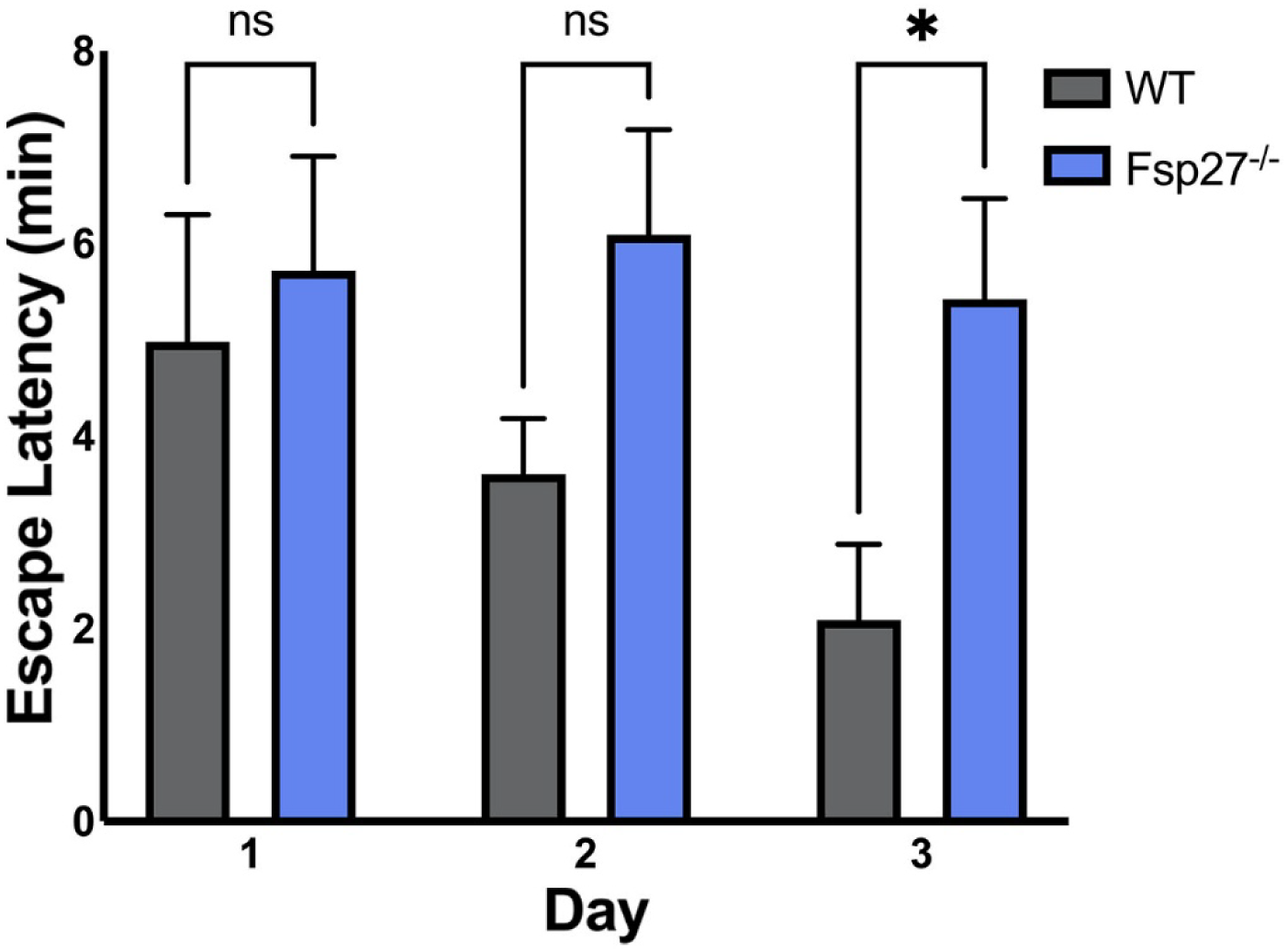
Labyrinth maze test in Fsp27^-/-^ and WT mice. Time taken to escape the labyrinth maze for WT and Fsp27^-/-^ mice across all three days. Data presented as mean ± SEM. *\*P* ≦ .05, *\*\*P* ≦ 0.01, *\*\*\*P* ≦ 0.001; Unpaired Student t-test.

### Y-Maze Reveals Compromised Spatial Working Memory in Fsp27^-/-^ Mice

To further study the cognitive impairment in Fsp27^-/-^ mice, we performed a Y-maze test, which is used for simple choice or exploratory behavior and spontaneous alternation in mice. The average percentage of spontaneous alternations was calculated for WT and Fsp27^-/-^ mice. Fsp27^-/-^ mice displayed a lower percentage of spontaneous alternations compared to WT mice indicating diminished spatial working memory, as they were unable to remember and choose different arms of the maze (Fig. 3A). Additionally, Fsp27^-/-^ mice made fewer total arm entries, suggesting a reduction in exploratory behavior and locomotor activity (Fig. 3B). The decreased alternation suggested hippocampal dysfunction and cognitive deficits associated with Fsp27 depletion. Also, a clear indication of decline in working memory, recognition memory and locomotor activity, the parameters associated with hippocampal function and short-term memory, was evident in Fsp27^-/-^ mice.

**Figure 3.**
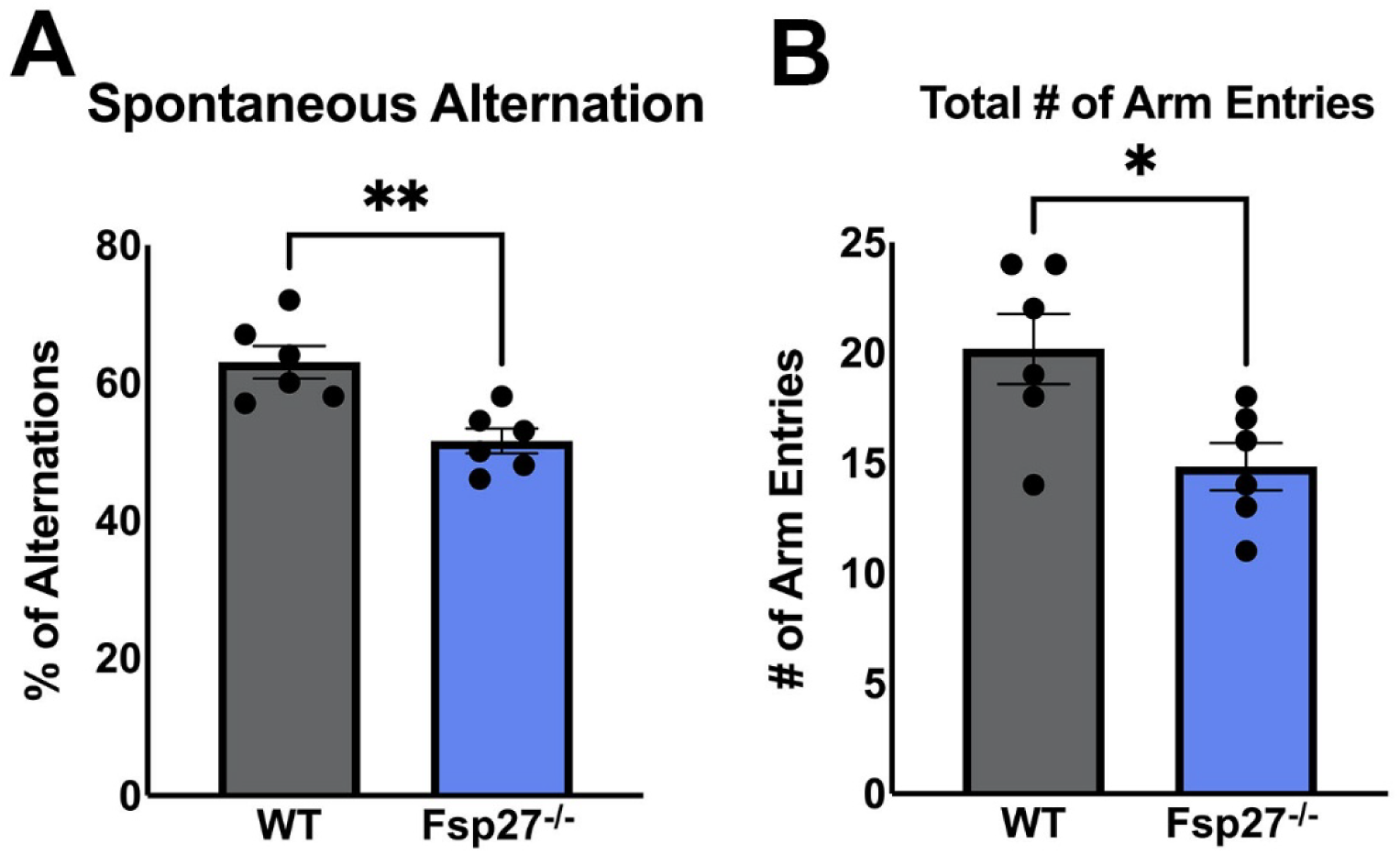
Y-maze test to assess hippocampal function and short-term memory in Fsp27^-/-^ mice. Time taken to escape the labyrinth maze for WT and Fsp27^-/-^ mice across all three days. Data presented as mean ± SEM. *\*P* ≦ .05, *\*\*P* ≦ 0.01, *\*\*\*P* ≦ 0.001; Unpaired Student t-test.

### RNA-Sequencing Reveals Gene Imbalance and Pathway Alterations in Fsp27^-/-^ Mice

To achieve insight into the molecular changes resulting from Fsp27 deletion, we conducted RNA-sequencing analysis on brain tissue samples from Fsp27^-/-^ and WT mice. After performing stringent quality control, normalization, and data pre-processing, differential expression analysis identified a total of 230 significantly differentially expressed transcripts (absolute fold change ≥ 1.5 and p-value < 0.05) between Fsp27^-/-^ and WT brain samples (Fig. 4A). Among these, 97 transcripts were significantly downregulated in the brains of Fsp27^-/-^ mice compared to WT mice, while the remaining genes were upregulated. The differentially expressed genes depicted differential expression patterns among Fsp27 knockout and WT samples as shown in the heatmap of the top 40 dysregulated genes (Fig. 4B). As expected, Fsp27/Cidec itself emerged as the most significantly downregulated gene in samples from Fsp27^-/-^ mice compared to the WT controls, validating the experimental design and transcriptomic data quality (Fig. 4A). In addition, several biologically relevant genes were among the top downregulated targets, including Rec8(36), a gene associated with genomic instability, Col6a4, involved in maintaining extracellular matrix integrity (37), and Lipg, a key regulator of lipid metabolism (38). Conversely, a subset of genes was significantly upregulated in the knockout condition, including Tnnt3, related to muscle function and contractility(36), Cyp2e1, a known driver of oxidative stress via reactive oxygen species (ROS) production(39), Adcy3 (Ac3), which mediates signal transduction through cAMP, and Ngp, a gene implicated in neuroinflammatory responses(40) (Fig. 4 A&B).

**Figure 4.**
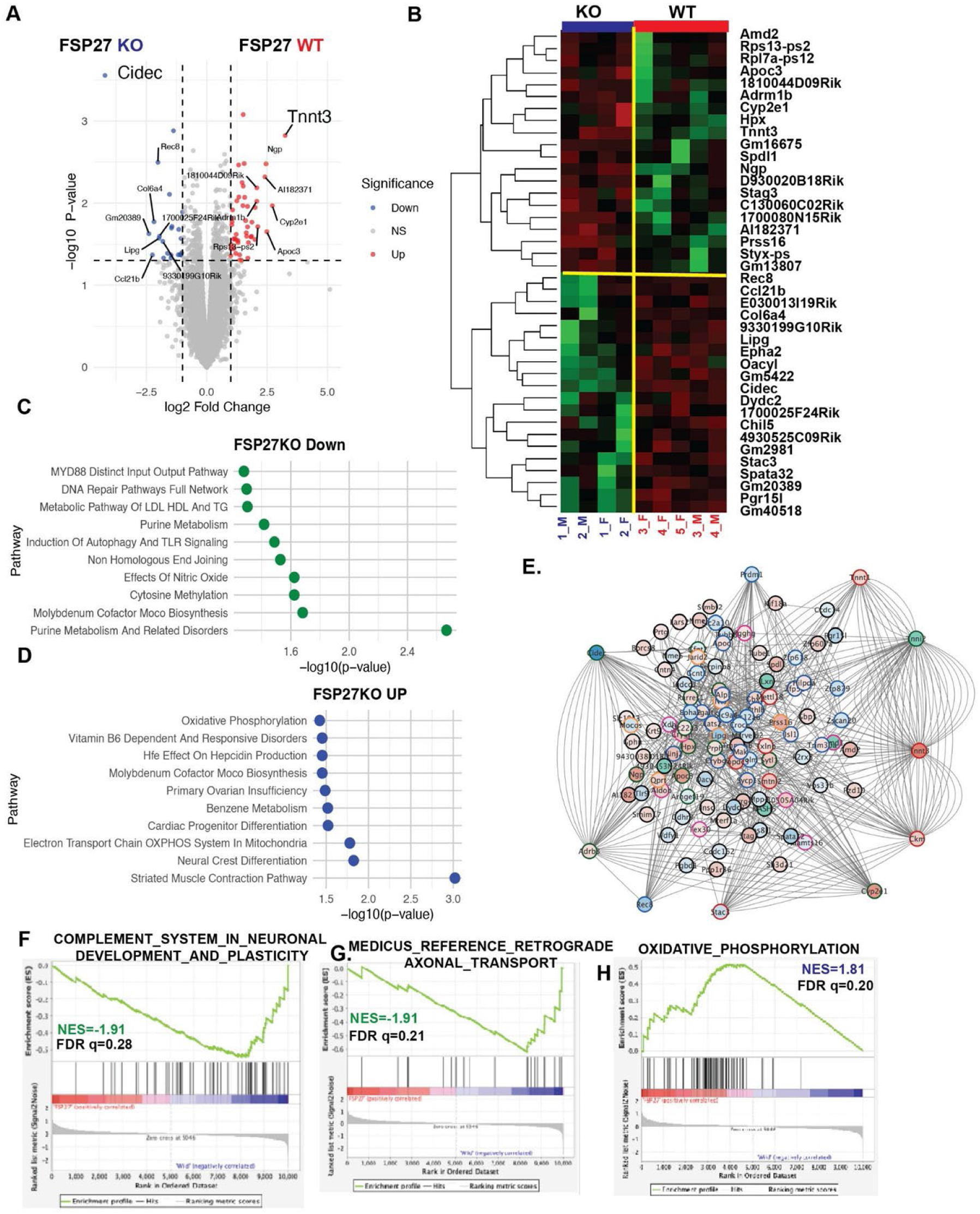
RNA sequencing depicts association of Fsp27 knockout with an enhancement of oxidative stress and suppression of neural and sensory processes A. Volcano plot showing differentially expressed genes between Fsp27^-/-^ (Fsp27 KO) and wild-type (WT) samples. Genes are colored by significance: upregulated (red), downregulated (blue), and not significant (grey). Key top genes are labeled. B Heatmap of top 40 differentially expressed genes between Fsp27^-/-^ mice and WT samples, illustrating expression patterns. In the heatmap, each row represents a gene, and each column represents a sample. Gene expression levels are shown using color: red indicates upregulated expression, while green indicates downregulated expression. C Pathway enrichment analysis of downregulated genes in Fsp27^-/-^ mice samples with selected top pathways dot plot based on raw P value (−log10(p-value) on x-axis). D. Pathway enrichment analysis of upregulated genes in Fsp27^-/-^ brain samples, showing top 10 enriched pathways based on raw P value (−log10(p-value) on x-axis). E. Interactive network illustrating genes that are significantly dysregulated in Fsp27^-/-^ mice samples compared to WT. In this network, each node represents a gene, and each edge indicates an interaction between genes identified using Genemania package in the Cytoscape platform. Nodes are colored according to fold change: red for upregulated genes and blue for downregulated genes. Master regulator genes, identified by the highest degree of connectivity, are positioned at the network periphery to highlight their regulatory roles. F–H. Gene Set Enrichment Analysis (GSEA) plots for the selected gene sets: Complement System in Neuronal Development and Plasticity (F), Axonal Transport (G), and Oxidative Phosphorylation (H).

Pathway enrichment analysis of the downregulated genes in Fsp27^-/-^ mice revealed significant (P <0.05) perturbations in purine metabolism, effects of nitric oxide, fatty acid metabolism, and DNA repair related pathways (Fig. 4C). In contrast, the upregulated genes were significantly enriched (P <0.05) in pathways related to oxidative phosphorylation, neuronal differentiation, and muscle contraction, suggesting a broader physiological shift in response to Fsp27 depletion (Fig. 4D). Additionally, interactive network analysis formed a cohesive network of genes which are up and down regulated due to Fsp27 depletion (Fig. 4E). The topological analysis on the interactive network identified Tnnt3 and Cyp2e1 as master regulators (based on degrees of interactions) previously linked to CNS function and oxidative stress, which may contribute to dementia, neurological diseases, and Alzheimer’s disease (Fig. 4E). Similarly, downregulated master regulators included Rec8, Prrdm1, and Stac3, which are critical for developmental processes encompassing chromosome segregation, neural and sensory processes, and muscle development. Downregulation of these genes may lead to developmental disorders, tissue dysfunction, or an increased risk of neuronal diseases. Additionally, gene set enrichment analysis (GSEA) also depicted that Fsp27 depletion is associated with negative enrichment of Neuronal Development and Plasticity (NES=-1.91, FDR q=0.28, Fig. 4F) and Axonal transport (NES=-1.91, FDR q=0.21, Fig. 4G) related gene sets that are directly associated with neuronal development and neurodegeneration related processes (41–43). Conversely, the FSP27 knockout was also associated with the upregulation of Oxidative phosphorylation (NES=1.81, FDR q=0.20, Fig. 4H) that might result in oxidative stress through increased generation of ROS driven by stressed or excessive mitochondrial respiration(44).

In summary, Fsp27 depletion may enhance oxidative stress and suppress neural and sensory processes, potentially reducing neural integrity and increasing the risk of future neurological diseases including dementia and Alzheimer’s.

## Discussion

Brain and skeletal muscle function have been observed to be interconnected through various pathways and mechanisms that involve both the brain and musculoskeletal system (45, 46). Our recent study suggested that FSP27 plays a significant role in the regulation of muscle function (26). Therefore, we hypothesized that FSP27 might play a role in cognitive function. Emerging studies have uncovered its involvement in hepatic and skeletal muscle function. Furthermore, despite growing insights into FSP27’s metabolic role in various tissues (12–21) , its involvement in the central nervous system and cognitive regulation have remained unexplored. Therefore, the present study was designed to specifically study the role of FSP27 in cognitive function in Fsp27^-/-^ mice. By assessing performance in maze tasks alongside other behavioral measures, we built a profile of the animal’s cognitive abilities and behavioral traits. Our findings contribute to a deeper understanding of the role of FSP27 in the processes linking cognition and metabolism. Behavioral assessments demonstrated that deletion of the Fsp27 gene impairs multiple domains of cognition, including learning, memory retention, and spatial recognition. In the Morris Water Maze, Fsp27^-/-^ mice exhibited delayed acquisition to the task, characterized by a slower learning curve and inefficient navigation strategies, suggesting deficits in learning and spatial recognition. Similarly, in the Labyrinth Maze, Fsp27^-/-^ mice demonstrated no improvement across trials, confirming an impairment in learning ability and cognitive flexibility. Furthermore, the Y-Maze test revealed a marked reduction in spontaneous alternation behavior among Fsp27^-/-^ mice, indicative of impaired spatial working memory. Collectively, these results suggest that Fsp27 plays a critical role in maintaining cognitive function, particularly in processes related to learning, memory retention, and spatial memory.

This study presents the first comprehensive transcriptomic analysis in Fsp27 depleted mouse brain, revealing broad alterations in gene expression associated with critical biological pathways. A total of 230 transcripts were significantly dysregulated in Fsp27^-/-^ mouse brain tissue, with a nearly balanced distribution of upregulated and downregulated genes, supporting a profound and multifaceted role of Fsp27 in central nervous system (CNS) homeostasis. Several genes known to regulate genomic stability (e.g., Rec8), extracellular matrix organization (Col6a4), and lipid metabolism (Lipg) were significantly downregulated in Fsp27^-/-^ brain, suggesting that Fsp27 loss may impact fundamental cellular and structural processes within the brain. Conversely, the upregulation of genes such as Tnnt3, Cyp2e1, Adcy3, and Ngp highlights a shift toward enhanced oxidative metabolism, altered contractility, and increased neuroinflammatory potential. Pathway enrichment analysis revealed significant perturbations in DNA repair, fatty acid metabolism, and nitric oxide-related signaling pathways among the downregulated genes, all of which are essential for maintaining neural integrity and cellular stress responses. The repression of these pathways may contribute to cellular vulnerability, particularly under metabolic or environmental stress. In contrast, oxidative phosphorylation, neuronal differentiation, and muscle contraction pathways were significantly enriched among the upregulated genes, possibly reflecting a compensatory or maladaptive response to disrupted energy and structural homeostasis.

Notably, interactive network analysis identified upregulated genes: Cyp2e1 and Tnnt3 as master regulators that are critical for the stability of gene network generated due to Fsp27 deletion. Cyp2e1 has been previously implicated in ROS production(39), oxidative stress(39), and neurodegenerative conditions, including Alzheimer’s disease and dementia (47, 48). This suggests that Fsp27 deficiency may lead to a sustained prooxidative state, which could compromise neuronal survival and synaptic plasticity over time. On the other hand, downregulated master regulators such as Rec8, Stac3, and Prdm1 are essential for chromosome segregation, neural development, and muscle integrity. Loss of expression of these genes has been associated with impaired developmental processes, tissue dysfunction, and increased risk of neurodevelopmental disorders(49–51). These findings align with GSEA results, which showed significant negative enrichment of gene sets related to Neuronal Development and Plasticity and Axonal Transport, which are two processes that are crucial for synaptic function, learning, and memory. Disruption of these pathways is a hallmark of several neurodegenerative diseases, strongly supporting a potential mechanistic link between Fsp27 loss and cognitive decline.

Future studies should explore whether restoring Fsp27 function or targeting key hub genes like Cyp2e1 could mitigate neurodegenerative processes and preserve cognitive function. Additionally, the role of Fsp27 in human neurological disorders warrants further investigation, particularly in the context of aging and metabolic diseases, where its regulatory effects on lipid metabolism and mitochondrial function may be especially relevant.

Previous studies have shown that the absence of Fsp27 is associated with increased mitochondrial oxidative metabolism (15). Despite this heightened metabolic activity, Fsp27^-/-^ mice exhibit reduced muscle performance and strength (26). Given the established role of muscle-brain communication in supporting cognitive functions(6, 7), these findings suggest a potential pathway by which Fsp27 may influence cognition, potentially through alterations in myokine signaling or energy availability. Additionally, the fat in the brain is vital for maintaining the structural integrity of neurons and supporting the myelin sheath that enables rapid nerve impulse conduction(4, 5). Lipids such as omega-3 fatty acids also play protective roles against oxidative stress, which is likely elevated in Fsp27^-/-^ mice and may contribute to neuronal cell damage (52). Thus, absence of Fsp27 in the brain may impair lipid homeostasis, compromising neurobiological processes crucial for learning, memory, and spatial awareness.

Several limitations of this study may have influenced the interpretation of the neurobehavioral test results. As noted previously, Fsp27^-/-^ mice exhibit reduced muscle performance and strength compared to WT mice(26). These physical deficits may have contributed to slower escape latencies in the Morris Water Maze and reduced performance in the Labyrinth Maze. However, both WT and Fsp27^-/-^ mice demonstrated comparable speeds during these tasks, suggesting that motor impairments alone do not fully account for the observed cognitive differences. Additionally, the Y-Maze test for spatial working memory requires minimal physical exertion, further supporting the conclusion that the impairments observed are cognitive in nature rather than purely motor-based.

Another limitation is the use of whole-body Fsp27 knockout mice, which makes it difficult to determine whether the effects on cognition are driven by the loss of Fsp27 in a specific tissue, such as adipose tissue, skeletal muscle, or the brain itself. Future studies utilizing tissue-specific knockouts would help clarify the site of action. This study was also limited to male mice, so further studies should examine any potential effects of sex differences on cognition.

In summary, our findings reveal a previously unidentified role of Fsp27 in regulating cognitive function. Loss of Fsp27 impairs learning, memory retention, and spatial working memory, as demonstrated across multiple neurobehavioral assessments. These cognitive impairments may stem from a combination of altered metabolic homeostasis or disrupted communication between the muscle and the brain. While this study provides important insight into the role of Fsp27 in cognitive function, further research is needed to identify the molecular pathways mediating these effects. Understanding how Fsp27 contributes to cognitive health may offer new therapeutic targets for metabolic and neurodegenerative diseases that involve cognitive dysfunction.

## Materials and Methods

### Mouse Maintenance, Generation, and Dissection

Fsp27^+/-^ mice were generously provided by Dr. Yoshikazu Tamori from Kobe University, Japan(15) and were of a C57BL/6N genetic background. Crossing of these heterozygous mice was performed to obtain homozygous Fsp27^-/-^ mice. For this study, male mice were used, all of which were 7-8 months of age during neurobehavioral testing and 9-10 months of age at the time of dissection. All mice were fed a standard chow diet and housed in the Ohio University Animal Facility (Athens, OH) with free access to food and water under a 12-hour light/dark cycle.

All experimental protocols were conducted in accordance with the Institutional Animal Care and Use Committee (IACUC) guidelines and were approved by the Ohio University IACUC. Ohio University holds an Animal Welfare Assurance (A3610-01) from the NIH-Office of Laboratory Animal Welfare and complies with the USDA Animal Welfare Act Regulations under license 31-R-0082. Care and use of laboratory animals adhered to the *Guide for the Care and Use of Laboratory Animals.* Carbon dioxide exposure was used to euthanize mice, followed by cervical dislocation to ensure humane treatment.

Tissues were immediately collected, snap-frozen in liquid nitrogen, and stored at −80°C for future molecular analyses.

### Morris Water Maze

The Morris Water Maze was conducted to evaluate spatial learning and memory. The experiment set-up consisted of a circular pool (180 cm in diameter, 40 cm deep) and was divided into four quadrants (Fig. 5A). The pool was filled with tap water and the temperature was maintained at 26 ± 1°C. A submerged escape platform (10 cm in diameter) was placed at a fixed location within quadrant 1 of the pool. Distinct visual cues were positioned around the testing room to aid spatial navigation. A camera was positioned above the pool to record trials for analysis.

**Figure 5.**
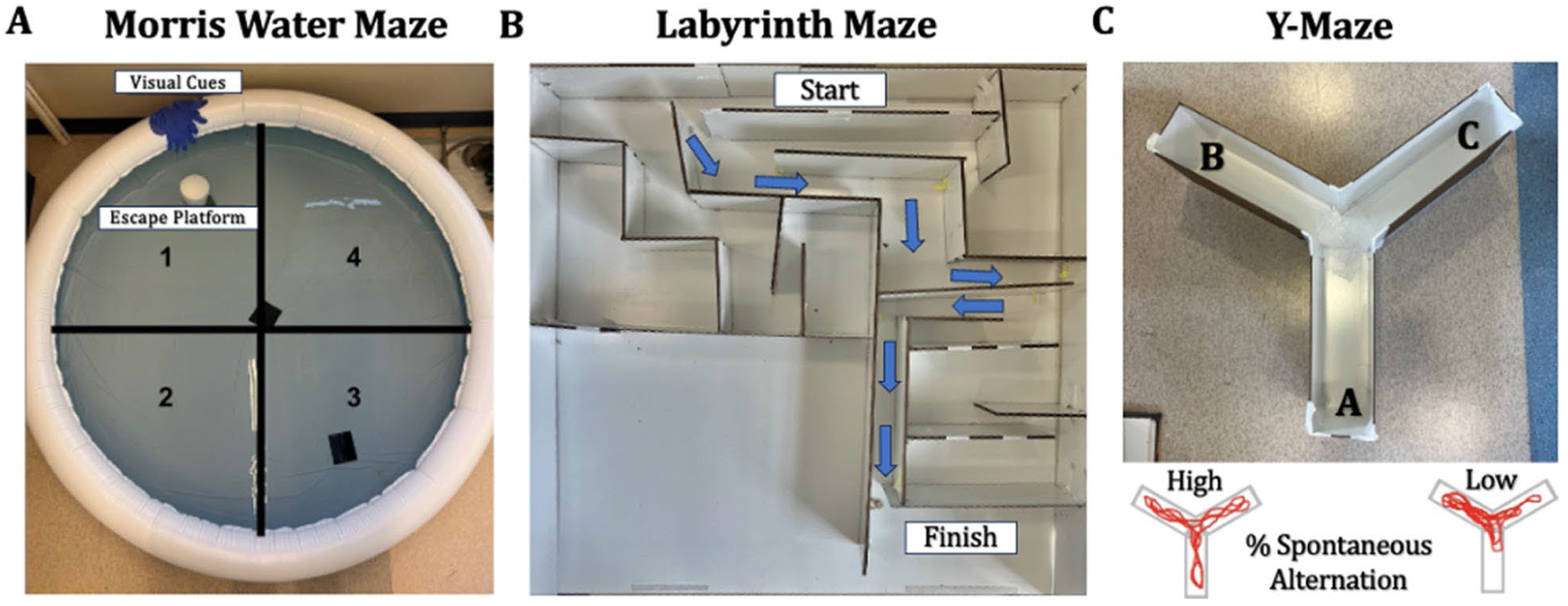
(A) The Morris Water Maze used in this study, divided into four quadrants as indicated by the black lines (not visible to the mice). (B) An image of the classic labyrinth maze used in this study with arrows indicating the correct path for reference. (C) Shows an image of the Y-maze used in this study with arms labeled A, B, and C for data analysis.

The water maze procedure was based on established methodologies in the literature and consisted of three phases: visible, training, and probe(32). Visible Phase (Day 1): The platform was placed in quadrant 1 of the pool and elevated 1 cm above the water’s surface. Each mouse was individually placed tail-end first, into quadrant 1 of the pool, facing the outer wall. Mice were allowed for up to 60 seconds to locate the platform and the time taken for each mouse to reach the platform (escape latency) was recorded. If a mouse failed to reach the platform within 60 seconds, it was gently guided to the platform and allowed to remain there for 10 seconds to reinforce its location as their escape from the pool. This process was carried out for all mice starting in quadrant 1, after which their starting point was shifted to quadrant 2, 3, and lastly 4. Between trials, mice were dried with a towel and returned to their home cage to rest.

Training Phase (Days 2-4): During the training phase, the platform was submerged 1 cm below the water’s surface in the same fixed location as the visible phase. The mice completed the same process and were allowed 60 seconds to reach the platform. The time taken for each mouse to reach the platform was recorded. Additionally, the path length to the platform for each mouse was recorded.

Probe Phase (Day 5): For the probe phase, the platform was removed from the pool, but the visual cues remained in place around the pool. Each mouse was placed in quadrant 1 of the pool and allowed 90 seconds to swim freely. The number of times a mouse crossed over the platform’s previous location (platform crossings) was recorded along with the time each mouse spent in the goal quadrant to measure spatial memory. This process was repeated with the mice starting in quadrants 2, 3, and 4.

### Classic Labyrinth Maze

The classic labyrinth maze was utilized to evaluate learning and memory in the mice (Fig. 5B) as described earlier (33). Before testing, mice underwent a habituation period of 60 minutes in the testing room to reduce potential anxiety-related behaviors. The experiment was conducted in an isolated area to minimize external interference.

On testing days, each mouse was placed at the designated starting point of the maze and allowed a maximum of 10 minutes to navigate to the exit or escape zone. A successful escape from the maze was defined by the mouse reaching the escape zone and remaining there for at least 10 seconds. The time taken for each mouse to escape the maze was recorded with a stopwatch. The apparatus was cleaned with 70 percent ethanol between trials to remove scent cues. This procedure was repeated on three consecutive days for a total of three trials.

### Y-Maze

The Y-maze was employed to assess spatial working memory measured by spontaneous alternation; a behavior driven by rodents’ innate curiosity to explore novel environments(34). The apparatus consisted of three opaque, light-colored arms, each oriented at 120° angles from each other (Fig. 5C). Each arm was labeled A, B, or C. Mice were acclimated to the testing room a week prior to the experiment to familiarize them with the environment.

The experiment included three trials in which the mice were placed in the distal end of arm A facing the center of the maze. The mice were allowed to explore the maze freely for 5 minutes and experimenters left the area during this period to avoid influencing the rodent’s behavior. The maze was cleaned with 70% ethanol and dried before testing each mouse. A camera was mounted above the maze to record each trial, and videos were saved for subsequent analysis.

Video recordings were reviewed to document the number of arm entries and alternations. An arm entry was defined by the mouse entering an arm with all four limbs. An alternation was defined as consecutive entries into all three arms (e.g., A → B → C). The percentage of alternations was calculated using the following formula: % Alternations = (Number of Alternations/Total Number of Arm Entries – 2) X 100. A higher percentage of alternations indicates better spatial working memory, reflecting the mouse’s ability to recall previously visited arms and avoid re-entry into the same arm (34, 35).

### Transcriptome profiling via RNA-sequencing to Investigate Molecular Changes After Fsp27 Deletion

To explore the molecular mechanisms underlying Fsp27 knockout, we performed high-throughput transcriptomic profiling by RNA sequencing on brain tissue of Fsp27^-/-^ mice (n=4) and WT (n=5) mice. Total RNA was extracted from both Fsp27^-/-^ and WT samples using the QIAGEN RNeasy Plus Mini Kit, which includes genomic DNA elimination steps to ensure RNA purity. The integrity and quality of the extracted RNA was confirmed using the Agilent Bioanalyzer 2100 and samples with high quality RNA (RIN>6) were processed further for RNA-seq libraries generation.

RNA-sequencing libraries were prepared using the Illumina TruSeq Stranded mRNA Library Prep Kit by capturing polyadenylated transcripts. The quality and fragment size distribution of the libraries were examined using the Agilent High Sensitivity DNA Kit. The high-quality libraries were sequenced on the Illumina NovaSeq 6000 platform, generating a minimum of 20 million paired-end reads per sample (PE150) to achieve deep and comprehensive coverage of the transcriptome. This depth enables reliable detection of both low- and high-abundance transcripts, facilitating differential expression and downstream pathway enrichment analysis to uncover the transcriptional impact of Fsp27 knockout.

#### Analysis of RNA-seq data

The demultiplexed raw FASTQ files were first subjected to quality control using FastQC to assess read quality, sequence duplication levels, GC content, and other key metrics. Adapter sequences and low-quality bases were subsequently removed using Trimmomatic to ensure high-quality reads for downstream analysis. Following trimming, the filtered high-quality reads were aligned to the mouse reference genome (GRCm38) using the HISAT2 aligner, a splice-aware aligner optimized for accurate mapping of RNA-seq data(53). Post-alignment, gene-level read counts were generated using HTSeq-count (54), which quantifies the number of reads mapped to each annotated transcript. This produced a raw count matrix representing transcript abundance across all samples, forming the basis for downstream differential expression and pathway analyses.

#### Normalization and differential Expression analysis

To identify transcriptomic changes associated with Fsp27 knockout, we first performed quality control and filtering of the raw transcript count data. Transcripts expressed in only one sample or with extremely low expression across all samples were excluded to reduce noise. Normalization was carried out using the VOOM algorithm(55) from the limma package, which adjusts for library complexity and sequencing depth while transforming count data to log2-counts per million (logCPM) for downstream linear modeling. The normalized data was used for differential gene expression analysis by implementing linear modeling and empirical Bayes moderation through the limma package (56). Genes were considered significantly differentially expressed if they had an absolute fold change ≥ 1.5 and an associated raw p-value < 0.05.

#### Pathways, Interactive network, and Gene Set Enrichment Analysis

To understand the biological relevance of the differentially expressed genes (DEGs), we performed pathway enrichment analysis using the Enrichr tool(57) with WikiPathways database(58). Pathways with a p-value < 0.05 were considered significantly altered due to Fsp27 deletion.

To explore the functional connectivity and gene-gene interactions, we conducted interactive network analysis using the GeneMANIA(59) package within the Cytoscape(60) platform. This analysis integrated curated gene-gene and pathway-level interactions from literature and public databases. The resulting network was analyzed to identify master regulators, defined as genes with the highest number of interactions within the network, suggesting their central role in mediating the transcriptomic effects of Fsp27 knockout.

In addition, to capture more subtle, coordinated changes at the gene-set level, we performed Gene Set Enrichment Analysis (GSEA) using the Molecular Signatures Database (MSigDB)(61) canonical pathways collection (C2 CP). Gene sets with an FDR q-value < 0.30 were considered significantly altered and reflective of biological geneset-level perturbations due to Fsp27 deletion.

## ACKNOWLEDGEMENTS

This work was supported by NIH grant R01DK101711 (VP), RO1DK138635 (VP), and funds from Osteopathic Heritage Foundation’s Vision 2020 to Heritage College of Osteopathic Medicine at Ohio University (VP).

## Competing Interest Statement

The authors declare no competing interests.

## Classification

Major: Biological Sciences, Minor: Neuroscience

## Notes

### Competing Interest Statement

The authors have declared no competing interest.

